# Perception/action coupling in children with autism: insights from looking time and pupil dilation measurements

**DOI:** 10.1101/2024.08.22.609181

**Authors:** Nicole Clavaud-Seon, Quentin Guillon, Judith Vergne, Sandrine Sonié, Christina Schmitz

## Abstract

The objective of this study was to characterize, through indices extracted from eye-tracking measurements, the spontaneous distinction of videos of daily actions with a variable perception/action coupling, depending on whether, for the same action, the video was presented in the forward reading direction (strong coupling), or in the backward reading direction (weaker coupling). 17 pairs of videos of daily actions performed by adults were viewed by 36 typically developing children and 28 children with ASD aged 7-18 years. During the exposure phase, they watched two videos of the same action (forward and backward) presented successively, before looking at these two videos in competition, in a second visual preference phase. During the exposure phase, all participants paid similar general attention to each of the videos. We found greater pupillary dilation for backward than forward actions in both groups, but significantly less in the ASD group. In the visual preference phase, both groups showed significantly greater looking times for backward actions over forward ones, with no difference between groups. If TD children perceived the kinematics of the backward videos as violating their expectations given the strong perception/action coupling they had already built over that action, on the contrary, the lower increased in pupil dilation found in ASD children could reflect altered perception/action coupling. This study confirms the validity of looking time and pupil dilation as behavioral and physiological markers that could be used in a 10-mn eye-tracking test to explore perception/action coupling in childhood and in ASD.

**Lay summary:** People with ASD often have difficulty understanding the actions of others. The fine understanding of actions requires a coupling between the action we observe, and its representation stored in our memory. This process might be challenged in autism. Here we used a 10-mn eye-tracking test to explore perception/action coupling in ASD. Participants were watching videos of daily actions. Looking time and pupil dilation were measured while participants watched videos of daily actions, and were found to be relevant indexes.

## Introduction

Autism Spectrum Disorder (ASD) is defined in the DSM-5 (American Psychiatric Association, 2013) as a neurodevelopmental disorder characterized by, on the one hand, qualitative deficits in communication and social interactions, and on the other hand by restricted behaviors, activities and interests. Although present from an early age, motor disorders in autism are not part of the diagnostic criteria, even though they could be considered as one of the first signs of autism (Harris, 2017; Papadopoulos et al., 2012; Rinehart & Mcginley, 2010; Teitelbaum et al., 2004). Furthermore, if motor disorders do not appear to be specific indicators of ASD, they clearly provide valuable information to the complex problem of early identification of ASD (Iverson et al., 2019). Disorders of motor adjustments were described as early as 1943, when Leo Kanner reported that ASD children do not adjust their posture in advance to the solicitations of a relative who holds out his arms to them. He also noted their awkwardness in motor performance, their failure to use non-verbal gestures to communicate with others, and repetition of identical actions. Since Léo Kanner’s description, numerous studies have highlighted the impact that motor atypicalities and impairments could have not only on fine and gross motor skills development of individuals with ASD (Cook et al., 2013; Cook, 2016; Whyatt & Craig, 2013), but also on daily functional abilities ranging from difficulty tying shoelaces, to difficulty using and understanding non-verbal gestures during social interactions (Attwood et al., 1988; Centelles et al, 2013). Hence, while not exclusive to autism, motor impairments and atypicalities may have a significant impact on how ASD individuals perceive and understand the actions performed by others and consequently have a transversal, almost insidious impact on communication and social interactions (Rinehart & McGinley, 2010).

An action can be considered as a coherent set of well-planned movement units oriented towards a goal, with varying degrees of finesse and complexity in the kinematics and with precise biomechanics that give coherence to the all from the beginning to the end. More precisely, what produces coherence to the perceived action is also the velocity in the execution of the movement, the grip strength, acceleration and jerk that are part of the kinematics. Motor learning and motor experience shape action since an early age and throughout life span.

Understanding someone’s action encompasses being able to predict the goal of that action. To this end, typically developed individuals use the relevant information provided by the kinematics of the movements of the person performing the action (Decroix & Kalenin, 2019; Stapel et al., 2012). Interestingly, infants own sensori-motor capabilities, and specifically their fine motor skills, determine their ability to discriminate impossible from possible human movements (Reid et al., 2005). Using eye-tracking, Morita et al. (2012) studied the perception of possible and impossible movements in typically developed children. They found that by the age of 12 months, infants already had a representation of the biomechanical constraints of the human body, such as how an arm can move within the limits of what is biologically possible. Like adults, infants spent more time looking at the elbow area during impossible arm movement. However, unlike adults, infants did not have pupillary dilatation as an emotional response to the unpleasant feeling generated by a biomechanically impossible arm movement.

Because of their motor atypicalities and impairments, it may be quite different for children and adults with ASD to recognize and understand the actions of others. However, many studies that have examined action goal understanding in ASD individuals have found it to be preserved (Falck-Ytter, 2010, 2012; Hamilton et al, 2007, 2009; Marsh & Hamilton, 2011: Tiziana-Zalla et al, 2009). Yet, it is not clear that they perceive the kinematics of a person performing an action in the same way as typically developing persons. Cook et al. (2013) investigated the kinematics of ASD movements by recording trajectory, velocity, acceleration and jerk while typical and ASD adult participants performed horizontal sinusoidal arm movements. All participants were also asked to categorize the observed movements as “natural” or “unnatural” in a biological motion perception task. ASD participants were found to move with an atypical kinematics. More, ASD participants biased perception of biological movements was correlated with the degree to which kinematics were atypical. Indeed, they over-classified biological motion as “unnatural”. Interestingly, this was also correlated with the severity of autism symptoms. Thus, in ASD, early motor impairments and atypical movement kinematics could disrupt the formation of action representations, affecting “the perception, prediction, and interpretation of the movements of others” (Cook, 2016), and potentially impairing the proper functioning of the action-perception coupling.

Indeed, motor simulation, also known as perception/action coupling, enables us to couple the observed action with the motor representations in memory thanks to the Mirror Neuron Mechanism (MNM) (Rizzolatti et al., 2009). This “motor resonance” would allow a fine and immediate understanding of non-verbal gestures, the goal of others actions and their intentions, also during social interaction observation. The MNM seems to be functioning at an early age (Gallese et al., 2009) and would be involved in imitation, empathy, theory of mind and language (Perkins et al., 2010). In ASD, many studies have shown a dysfunction of this mechanism in adults (Bernier, et al., 2007; Hadjikhani, 2007; Hadjikhani et al., 2006 Honaga et al., 2010; Oberman et al., 2013) and in children (Dapretto et al., 2006), while other studies do not find such a dysfunction (Falck-Ytter, 2010; Fan et al., 2010; Marsh & Hamilton, 2011; Dumas et al., 2014; Poulin-Lord et al., 2014). Beyond this dichotomy, studies have shown that the perception/action coupling can be influenced, i.e. modulated, by different factors such as familiarity with the observed action (Calvo-Merino et al., 2006), to the motor experience, and even more to the motor expertise (Calvo-Merino et al., 2005; van Elk et al., 2008).

To study the coupling between perception and action, many studies have measured brain activity using EEG or fMRI. Much less invasive, eye-tracking allows the recording of behavioral and physiological markers of visual exploration that are eye movements and pupil dilation. Eye movements, through saccades and fixations, reflect the orientation of explicit visuo-spatial attention (Jeannerod et al., 1968). Pupil dilation is linked to the activation of the sympathetic system leading to the stimulation of the iris dilator muscles, and more precisely by the activity in the Locus coeuruleus Norepinephrine (LC-NE) system (Wang & Munoz, 2015). Pupil dilation occurs when there is a variation in cognitive load resulting from the processing of information that recruits a more or less important part of our resources (Papesh & Goldinger, 2012), or can be induced by emotional arousal (Sirois & Brisson, 2014). Indeed, an Event Related Pupil Dilation (ERPD) can translate an emotional arousing such as pain when viewing videos of an elbow bent backward (Morita et al., 2012) or surprise when an image of an animal (a dog) presented on a screen does not match the sound (neighing) of it (Krüger et al., 2020).

Here we used an eye-tracking paradigm to extract, in an ecological and non-invasive way, behavioral and physiological cues reflecting the modulation of the perception/action coupling in typically developed (TD) children and in ASD children when viewing videos of daily actions. Our main objective was to characterize, through visual exploration, looking times and variations in pupil diameter, the spontaneous distinction of daily actions presenting with a variable perception/action coupling depending on whether the video was presented in the forward reading direction (i.e. allowing a strong coupling between the observed action and the action that the participant can perform), or the same video presented in the backward reading direction (in this case, the coupling would be weaker).

We expected to find in typically developing children a greater pupillary dilatation for the backward videos, as compared to forward videos. This increased attention for backward videos was hypothesized because they differ from the kinematics of the actions that belong to their repertory. We anticipated this difference to be less marked in the ASD group because of their altered motricity. Further, during a visual preference test, we assumed that if children were sensitive to the action kinematics, this should generate an increased attention to backward videos, as characterized by increased looking times for the latter compared to forward videos. As for pupil dilation, we expected the preference for backward videos to be significantly greater in the control group than in the ASD group.

## Methods

### Participants

Participants with ASD were 28 (1 female, 27 males), between the ages of 7.1 and 17.8 years. They were recruited from specialized care centers, local parent advocacy groups and the local Rhône-Alpes Autism Resources Center. All participants had received prior to the study a clinical diagnosis of ASD according to the DSM 5 criteria. Their diagnosis was confirmed by a child psychiatrist based on the administration of the Autism Diagnostic Interview-Revised (ADI-R; Lord et al., 1995) and/or the Autism Diagnostic Observation Schedule (ADOS-G or ADOS-2; Lord et al., 2000). Perceptual Reasoning was assessed with the Wechsler Intelligence Scale for Children (WISC-IV) in 23 participants with a mean score of 102.7 (SD = 20.3, range 69 to 150). Perceptual Reasoning could not be assessed in 5 participants (one was excluded after the pre-processing stage).

The non-autistic, typically developing, comparison group consisted of 36 participants (20 females, 16 males) between the ages of 7.1 and 17.3 years, all attending regular school. The groups were matched with respect to their chronological age. Exclusion criteria were any known associated genetic, neurological or psychiatric disorders, seizures, visual impairments or medications that may affect pupil dilation. The study protocol was approved by the Ethics Committee, CPP Sud-Est VI (Ref: AU 177; Ref ID-RCB: 2015-A00-193-4). All children fully consented to participate and written informed consent was obtained from parents or legal guardians prior to the study.

### Stimuli

Seventeen videos of daily actions (as lace up a shoe, fold a T-shirt, put on a jacket…) were created. Videos were shot in a neutral room with constant ambient light set at 100 lux and depicted an actor or an actress dressed in neutral clothing. They were filmed from a three-quarters point of view so that the actions were clearly visible to the participants. The start and end positions of the actors were similar and also kept constant across actions so that regardless of the direction of the video (forward or backward) the start and end positions were the same. The sound of videos was removed and they were converted in grey scale and set to 408×720 pixels in resolution.

The 17 couple of forward and backward videos depicted daily actions with a clear goal, beginning and end, and with little effect of gravity on backward videos. Indeed, the effect of gravity on the actor’s clothes, or linked to the object used by the actor could constitute a bias because it could give the action displayed in the backward direction an almost impossible and weird side and not mainly due to abnormal kinematics. Importantly, for each action, the backward and forward videos were similar in terms of amount of information, movement, duration and brightness.

### Eye tracking

Pupil size and eye movement data were measured using an EyeLink 1000 eye-tracker (SR Research Ltd, Canada) in a free-head mode and at a sampling rate of 500 Hz (monocular, dominant eye). Participants sat in a comfortable armchair at a distance of 60 cm from a 20’’ display screen (1024×768 pixels). The experiment took place in a soundproofed room with indirect lighting kept constant across participants (illuminance level ∼ 10 lux). The experiment was implemented using Experiment Builder (SR Research Ltd, Canada).

### Procedure

The experimenter first explained to the participant how the session was to be conducted. Prior to the test phase, two samples of daily action video were presented and commented by the experimenter to illustrate what forward and backward actions meant. The instruction was to watch the videos of actions carefully, as if participants were at home quietly watching television. Subsequently, a nine-point calibration was performed in two times where participants were first required to follow and fix a black dot (target) that moved randomly in nine locations on the screen. The grid of the first calibration phase was definitively accepted if the second calibration phase indicated an average deviation of less than 1.5° between the recorded targets and eye positions. Once the calibration phase was completed, a central point was displayed before the first trial started, which could not begin until the experimenter ensured that the participant had carefully fixed this center point. Following the calibration, 17 videos of daily actions were presented to participants in a random order, excepted for the first trial, which was the same for all participants. This first trial was discarded from the analysis.

A trial consisted of an exposure phase followed by a preference phase (Fig. 1). In the exposure phase, a grey screen was first shown to participants for 2000 ms followed by a 500 ms central fixation point. Next, either on the left or the right side of the screen, an image of the first frame of an action was presented for 2000 ms, followed by the video for 5000 ms. Afterwards, the same sequence of events was repeated with the same action but played in the other direction and on the opposite side of the screen. In the preference phase, a 2000 ms blank grey screen followed by a 500 ms central fixation point were first shown to participants followed by the images and videos from the exposure phase displayed side by side on the screen. The order of the 17 couple of forward and backward actions was randomized across participants. Furthermore, the presentation side as well as action directions were counterbalanced across trials. The whole process lasted for about 10 mn.

**Figure 1.**
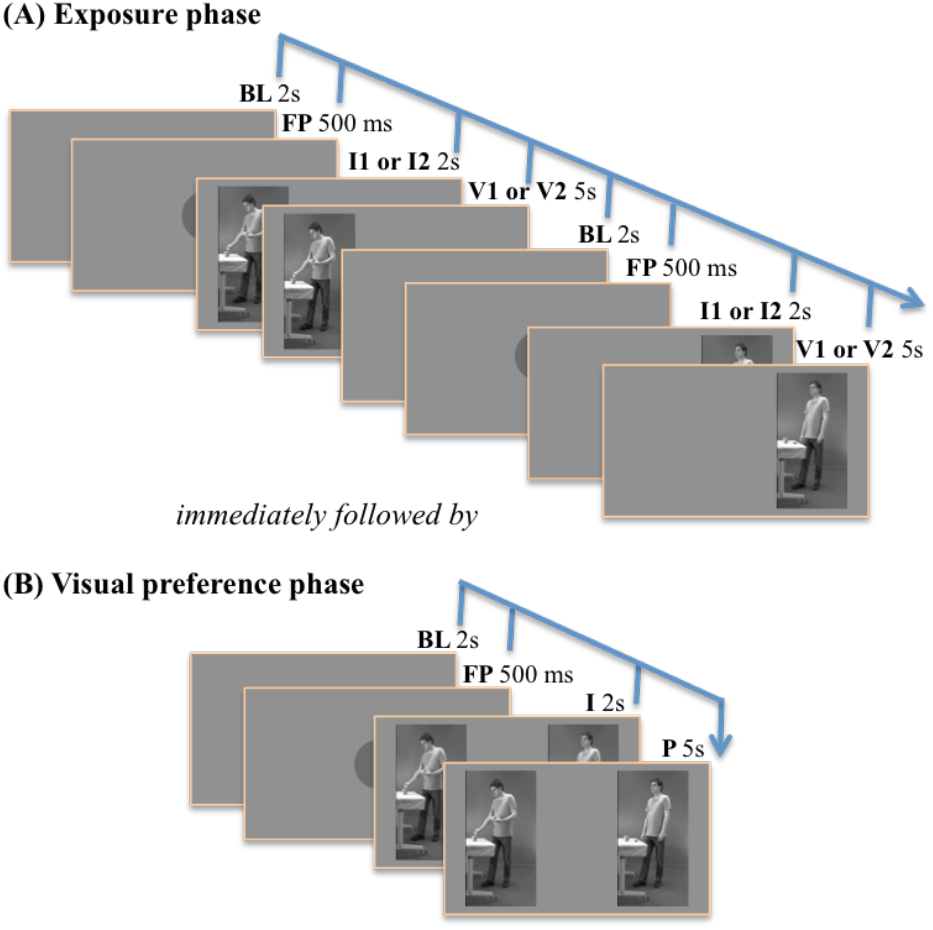
Experimental design showing the temporal sequence of the Exposure phase (**A**) and the Visual preference phase (**B**). BL: Baseline; FP: Fixation Point; I: Image; V: Video; P: Presentation.

### Pre-processing of pupil size and eye movement data

All pre processing stages were performed using Matlab (R2014). Only trials with less than 30% of missing data were kept for further analysis, resulting in the removal of 21 (2.2%) trials out of a total of 944. Inspection of the number of trials by participant indicated that one participant in the ASD group only had 5 trials (out of 16) satisfying this cutoff. This participant was removed from further analyses.

Preprocessing started with the identification and removal of blinks and blink-related artifacts (opening and closing periods of the eyelid). A pupillometry noise-based algorithm (Hershman et al., 2018) was used as well as angular velocity data computed from the x and y coordinates after the application of a Savitzky-Golay filter (Nyström & Holmqvist, 2010). A maximum velocity of 1000°/sec was used to detect periods of unphysiological eye movements, and an adaptive velocity threshold computed based on the average noise level, was then applied to determine the beginning and end of the period.

#### Pupil size

Blink and blink-related artifacts were first linearly interpolated. After interpolation, pupil data were smoothed using a zero-phase low-pass Butterworth filter (4Hz) to remove fast instrument noise, and down sampled from 500 to 20 Hz by taking the median pupil size per time in (50ms) to reduce computational cost. Pupil data were then aligned to the onset of each action video and segments of interest were created by extracting data between -200 pre- and +4820 ms post-video onset. Baseline correction (average pupil size in the time window from -200 to 200 ms) was applied using a subtractive method on a segment-by-segment basis. Segments where blinks and blink-related artifacts exceeded 30% of the total segment sample were rejected from further analysis. A total of 1798 segments were retained (out of 1858). The average number of segments per participant was 31 (range 24 to 32) and 30 (range 24 to 32) for the control and ASD group respectively. No further participants were excluded. For each participant, an average pupil size change was computed for backward and forward actions with a winsorized mean.

#### Eye movements

Total looking time towards either backward or forward actions during the exposition phase was computed by multiplying the total number of gaze points falling onto the respective action video by the sampling time interval (i.e. 2 ms). For the preference phase, preference for backward relative to forward actions was computed by dividing the number of gaze points that fell onto the backward action by the total number of gaze points that fell onto any actions (backward + forward). Thus, the preference represents the probability of looking at backward actions, given that the participant is looking at an action, whether backward or forward, and range from 0 to 1. Trials were excluded if 1) one of the two actions was not looked at (defined as less than 100 ms of looking time), either backward (n=64) or forward (n=101), and 2) the percentage of blink or blink-related artifact samples exceed 30% (N=12, nTYP=3; nASD=9). This last criteria was decided upon inspection of the distribution of the number of missing gaze data points across the whole sample. As a result, a total of 749 trials (79%) were analyzed (out of 944). The median numbers of trials per participants out of 16 was 14 (range 7 to 16) and 13 (range 8 to 16) for the control and ASD group respectively. No further participant was excluded.

### Statistical analyses

We used R (Version 3.6.1; R Core Team 2019) and the R-packages *lme4* (Bates et al. 2015), *lmerTest* (Kuznetsova et al., 2017), *tidyverse* (Wickham et al., 2019), *WRS2* (Mair & Wilcox, 2020) for data preparation, analysis and presentation.

#### Curve shape of pupil change

We performed a growth curve analysis of the pupil size (Mirman et al., 2008). Growth curve analysis is a multi-level modeling technique that allows the changes in pupil size to be modeled with orthogonal polynomials representing key aspects of the curve shape. For the current study, the pupil time course was modeled with a third-order polynomial and fixed effects of group (ASD vs. control) and direction (backward vs. forward) and their interaction on all time terms. In this model, the intercept refers to the mean pupil size over the full segment; the linear term refers to the slope of the pupil time course, with lager values indicating lager pupil size at the end than at the beginning of the segment; the quadratic term refers to the shape of primary inflection, with more positive values indicating a flatter inverted-U shape and more negative values, a steeper inverted-U shape; and the cubic term refers to the extent to which there is a secondary inflection, with positive values indicating an earlier shifted peak and negative values indicating a later shifted peak response (Kuchinsky et al., 2013). The random structure included participant random effects and participant-by-condition random effects on all time terms (i.e. intercept varying among participants and among directions within participants). GCA were fitted with maximum likelihood estimation and parameter estimates, degrees of freedom and corresponding p-values were estimated using Satterthwaite’s method. (i.e. random slope of direction within participant with correlated intercept).

#### Gaze behavior

Given the bounded nature of the preference score and the by-experimental-design dependencies evidenced at both the level of participants (each participants contributed multiple actions) and actions (each actions is viewed by multiple participants), generalized linear mixed effect models (GLMMs) with cross random effects for participants and actions were used (Jaeger, 2008). GLMMs were fitted with maximum likelihood estimation assuming a binomial conditional distribution and logit link function. The maximal random effect structure that is random intercepts for participants and actions and random slope for group within actions with correlated intercept was used (Barr et al., 2013). Group (ASD vs. TYP) was entered as a fixed effect and coded as a dummy variable (TYP = 0; ASD = 1) with TYP as the reference level, such that the model intercept represented the estimate for the TYP group and the beta coefficient represented the change in preference towards backward actions as a function of group on the log-odds scale (i.e. the estimated difference between TYP and ASD accounting for by-participant and by-action variation). Odds Ratio (OR) and their 95% bootstrapped CI are also reported for ease of interpretation. Parametric bootstrap was used to test the statistical significance of the fixed effect of group. Model comparison was also carried out to test the statistical significance of the fixed effect of group comparing the change in residual deviance between the full model and a model without the fixed effect of group using a likelihood ratio test. The simulation-based approach provided by the DHARMa package for residuals checks of GLMMs revealed overdispersion. An observation-level random effect (OLRE) approach was used to account for this. As a result, the random slope for group within actions was removed as non-convergence / singularity was detected.

## Results

### Pupil variation analyses

#### Control group

A significant difference in pupil terms between backward and forward actions was found for the intercept (*Estimate* = 436.50, *ES* = 7.77, *p* < 0.001), the linear (*Estimate* = 264.12, *SE* = 43.00, *p* <.001), and the cubic terms (*Estimate* = -67.96, *SE* = 16.13, *p* <.001), indicating that 1) the average pupil size was larger for backward actions (intercept), 2) the pupillary response increased more for backward than for forward actions (linear term) and 3) peaked at a later time (cubic term).

#### ASD group

A significant difference in pupil terms between backward and forward actions was found for the cubic term (*Estimate* = -86.81 *SE* = 29.91, *p* <.01), indicating that the pupillary response peaked at a later time for backward than for forward actions.

#### Comparison between the control and ASD groups

The analysis revealed a group by direction significant interaction for the linear term (*Estimate* = -145.09, *SE* = 72.35, *p* = 0.049), indicating that the backward pupil dilation showed a greater dilatation slope relative to forward actions in the control group compared to the ASD group. All other interaction terms did not reach statistical significance with the current sample size. A further investigation of the simple effect of group across directions on the linear term revealed a group difference for backward action (*Estimate* = -222.71, *SE* = 106.79, *p* = 0.041) and no group difference for forward action (*Estimate =* -77.63, *SE* = 91.49, *p* = 0.400), indicating that the interaction was mainly driven by a group difference on backward actions (Fig. 2).

**Figure 2.**
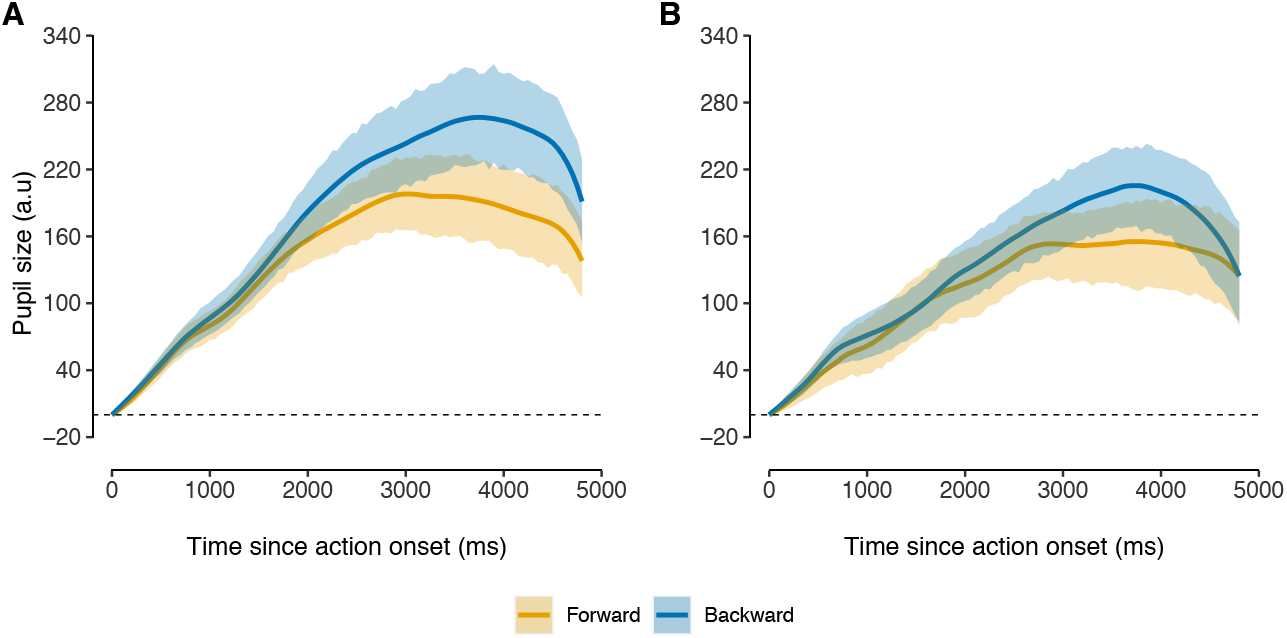
Time course of the pupil size for Forward and Backward videos in the control group (**A**) and in the ASD group (**B**). Solid lines represent the observed pupil size and shaded areas the 95% bootstrapped confidence intervals.

### Gaze behavior analysis

A simple effect of direction, as evidenced by a preference towards backward over forward videos was found in both groups independently. Indeed, accounting for the by-participants and by-actions variation, the odds of looking at backward videos relative to forward videos in the ASD group was 1.23 with a 0.95 bootstrapped confidence interval of [1.02, 1.49], and for the TYP group was 1.53 with a 0,95 bootstrapped confidence interval of [1.29, 1.78]. This indicated that all plausible values were greater than 1.00 in both groups and were therefore consistent with the rejection of the null hypothesis that there was no preference for backward over forward videos.

We did not find evidence against the null hypothesis that the ASD and TYP groups showed similar gaze patterns towards backward relative to forward videos with the current sample size (*Estimate* = -0.22, *SE* = 0.12, *p*_*boot*_ = 0.087). The odds of looking at backward videos relative to forward videos in the ASD group was estimated to decrease by a factor of 0.80 in comparison to that estimated for the TYP group. The 0.95 bootstrapped confidence interval of [0.63, 1.02] indicated a wide range of plausible values, consistent with a decreased of the odds of 0.63 and all the way up to an increased of the odds of 1.02 for the ASD group in comparison to the TYP group.

#### Total looking time during the exposure phase

Total looking time towards backward and forward videos were averaged (winsorized mean) across actions by participant. For backward videos, the average looking time was 4632 ms (min = 4124, max = 4769) and 4580 ms (min = 4100, max = 4771) for the TYP and ASD groups respectively. For forward videos, the average median looking time was 4670 ms (min = 4322, max = 4775) and 4604 ms (min = 3913, max = 4775) for the TYP and ASD group respectively. The median of all pairwise differences for backward videos was 42.3 ms with a 0.95 confidence interval of [-27.5, 116.1] and for forward videos, 50.0 ms with a 0.95 confidence interval of [-11.1, 112.1].

#### Total looking time during the preference phase

The average total looking time towards videos (backward and forward) was 4359 ms (min = 3584, max = 4519) and 4281 ms (min = 3564, max = 4534) for the TYP and ASD group respectively. The median of all pairwise differences was 68.8 ms with a 0.95 confidence interval of [-16.3, 186.7]. Total looking time during the preference phase was thus of the same order of magnitude between groups.

#### Spatial distribution of gaze during the preference phase

Kernel density maps for each video (forward and backward) by group (TYP and ASD) were computed from the raw gaze data point, with a smoothing bandwidth equal to 1°, and illustrate the distributions of gaze location across actions. Both groups tend to focus on the same features across actions, as revealed by visual inspection (see Supplementary material 1).

#### Excluded trials during the preference phase

Due to the limitation of the logistic regression when p = 0 and p = 1 (or p close to 0 or close to 1), trials where only one video was looked at by participants (operationalized as less than 100 ms of looking time) were excluded from the main analyses (see methods). Overall, 165 trials (17%) had to be removed, including 64 backward and 101 forward trials. A poisson mixed regression was used to model count data of trials where only one video was looked at. The random effect structure included random intercept for participants and a random slope for direction within participants. Group (TYP = 0, ASD = 1), direction (backward = 0, forward = 1) and the group by direction interaction were entered as fixed effects. Parametric bootstrap (n=1000) was used to test the statistical significance. Residuals checks, using the DHARMa package did not reveal any problem of uniformity nor dispersion. We did not find evidence against the null hypothesis that the effect of direction was similar across groups with the current sample size (*Estimate* = -0.76, *SE* = 0.42, *p*_*boot*_ = 0.065). The probability of looking only towards forward videos increased from a factor of 2.9 with a 0.95 confidence interval of [1.53, 6.8], meaning that across groups, forward videos tended to be more likely exclusively looked compared to backward videos.

## Discussion

Using eye-tacking, we designed a paradigm composed, first, of an exposure phase where participants separately visualized two videos of the same action presented in the forward reading direction, or in the backward reading direction. This was followed by a visual preference phase in which participants visualized the two videos side by side, in competition. We investigated, through the measurement of pupillary diameter and looking times, the spontaneous distinction of everyday actions presenting with a variable perception/action coupling depending on whether the action was presented in the forward reading direction (strong coupling), or in the backward reading direction (weaker coupling).

The exposure phase enabled to verify that every child had shown enough attention to both videos, in order to be able to interpret the visual preference test. First, total looking time did not differ for both directions (forward and backward) and was of the same order of magnitude between groups, thus limiting the possibility that the difference observed in the main analyses was a result of a difference in attention towards videos in the exposure phase. This further indicates that all participants had already seen both videos before the visual preference phase, thus, during the second phase, participants really asserted their preference for one of the two. In TD children, we found, as expected, greater pupillary dilation for backward videos, revealing increased attention compared with forward videos. The same was found in ASD children, but significantly less marked than in TD children. Moreover, during the visual preference phase, both groups showed an overall preference towards backward videos compared to forward videos, characterized by greater looking times.

Increased pupillary dilation and preference for the backward videos might be argued to be due to their weirdness, or their improbability. Nevertheless, more than half of the daily actions were as plausible in the forward as in the backward directions (e.g. the action in the forward direction “pull the zipper of a jacket down” becomes in the backward direction “pull the zipper of a jacket up”. They could both correspond to the participants’ action representations with the only exception that these two videos differ in their dynamics and kinematics, that is to say in what composes an action and gives its coherence: the velocity in the execution of the movement, the grip strength, acceleration, jerk and the precision level of fine motor gestures. If, when displayed backward, the video produced kinematic cues that made the whole action look “weird”, we assume that this was because the action no longer corresponded exactly to the observer’s motor representations.

The great pupillary dilation for backward videos found in TD participants in the exposure phase might thus reflect the surprise generated by differences in the kinematics and in the general coherence of the observed action. This is in line with Morita et al (2012), who found an increased pupillary dilatation in a group of adults, which would reflect the emotional response to viewing such an unpleasant movement that is an elbow bent backwards. Infants aged 6 and 12 months also showed increased pupil dilation when viewing an unexpected change when they looked at non-rational actions (moving food-laden spoon to the hand rather than mouth) (Gredebäck & Melinder, 2010). Interestingly, pupillometry could also be employed to detect surprise induced by violation of an expectation across a wide age range (Krüger et al., 2020), which is consistent with our findings. Variation of pupil size is known to be an indirect index of the LC-NE system activity (Murphy et al, 2014; Gilzenrat et al, 2010), which relies on two modes: a tonic activity mode that corresponds to the overall level of arousal and behavioral flexibility, and a phasic activity mode, in response to stimuli saliency (Aguillon_Hernandez, 2020; Devilbliss & Waterhouse, 2010; Sales et al, 2019). More specifically, in phasic mode, Event-Related Pupil Dilation (ERDP) may reflect the level of emotional arousal in response to surprise or violation of expectations, or errors in judgment of uncertainty following stimulus presentation (Krüger et al, 2020; Preuschoff et al., 2011, Sales et al., 2019).

Looking time comparison between the two directions of actions during the visual preference phase reinforces our pupillometry findings, as TD children asserted their preference for backward videos. Having already seen both videos during the exposure phase, our participants probably had no difficulty identifying the purpose of the videos presented, and could choose to spend more time watching the videos backwards. Identifying an action’s goal and the actor’s intention, based on its kinematics, are necessary to understand an action. Decroix et Kalénine (2019) explained that the attention of the observer would be first captured by the prediction of the goal of the actor’s action and next the observer would use the relevant information provided by the kinematics to confirm his prediction. Interestingly, already at 12 months infants observing actions focused on goals in the same way as adults did (Falck-Ytter et al., 2006). Given their age, it is highly probable that our TD participants relied on the kinematics cues of the actor movements to understand that one of the two videos was in a backward direction or at least different. Further, given that all TD participants viewed videos of everyday actions related to the use of objects they knew and used themselves (see list of actions in Supplementary material 1), we assume that they had most likely constructed representations of these actions based on their rich and well-developed motor skills, fueled by regular motor experience with these actions. Hence, TD children might have looked longer at the backward videos because their own motor skill was sufficiently advanced so that they perceived the kinematics of the backward videos as violating their expectations given the strong perception/action coupling they had already built over that action.

In the exposure phase ASD children also showed a significant increased pupillary response to backward videos as compared to the forward ones. However, this pupillary response was significantly less marked in the ASD group than in the TD one. The difference between the two groups cannot be explained either by the quality of the data nor by a difference in exploration of daily action videos during the exposure phase. Conversely, because of their locally oriented perception (Mottron et al., 2003) children with ASD might have been expected to detect inconsistencies in backward video more readily than TD children, resulting in greater pupil dilation for these videos in the former than in the latter. However, we found the opposite pattern. Although the way children with ASD explored each video was close to that of typical children (see kernel maps in Supplementary material 1), there is no guarantee that they captured the same cues as TD children to understand that backward videos were different from the forward one. In addition, the stimuli used in this study were not objects that could have triggered a specific interest for ASD participants, but dynamic daily actions, performed by real adult actors.

ASD participants not only needed to identify the goal of the action, but also to pay attention to the biological movements of the actors performing actions in order to be able to process the kinematics and pick up cues to understand finely that the video was backward. This process may have been undermined here, as it has been shown that people with ASD seem to rely more on the functional information given by objects, rather than the intentional information provided by the agents performing an action in order to understand it (Boria et al., 2009). Moreover, the fact that ASD individual had difficulties to perceive and categorized correctly natural movement of others have been suggested to be caused by their difficulty to pay attention to biological movement and to their atypical motor skills (Cook, 2016; Cook et al., 2013). Again, ASD participants’ sensitivity to the details of the actor’s kinematics may be lower than that of TD participants, as their motor and action representations may be different from those of TD children, and could distort their perception of others’ movement. Given their increased pupillary response to backward videos compared to forward, ASD children in this study showed that they had some kind of representation of the actions they watched, given also their own experience of these daily actions. But the lower increased in pupil dilation found in ASD compared to TD children could reflect that ASD participants may not see, or may not have access in the same way as typical participants to the fine details of the actor’s kinematics performing the action. This might have impacted the perception/action coupling strength in ASD participants, hence producing less violation of their expectations when viewing the backward videos for the first time.

Conversely, if during the exposure phase ASD children showed less pupillary dilation for backward videos, no difference could be found between the two groups concerning the looking times at backward videos during the visual preference phase. Interestingly, Somogiy et al. (2013) and Hamilton et al. (2007) showed that ASD children, including those with an intellectual disability, could recognize and imitate actions when they had a clear goal. Here, all actions had a clear goal, a clear beginning and a clear end. Marsh et al. (2015) in their eye-tracking study, which aimed to test whether gaze behavior during observation of an action was modulated by the rationality of an action, showed that basic goal understanding was preserved when the attention of ASD participants was maintained and they paid attention to salient features of the action. In our study, the instruction given to our participants was to carefully look at the actions performed by the actors. It is also worth noting that we were very careful that the actors displayed a minimum of social cues: they never looked at the camera, thus avoiding any direct eye contact, their faces were neutral and their movements were precise and clear, hence facilitating attention raised to the videos, and most probably action understanding. Finally, because the videos were presented for the second time during the preference phase, the increased looking time measured during the 5000 ms duration of the videos may have given them an advantage in capturing the difference in kinematics that ultimately directed their attention more toward the backward than the forward videos.

The fact that we did not evaluate ASD participant’s motor skills, nor did we evaluate their experience for the everyday actions they visualized constitutes a limitation of this study. For instance, we could have them reproduce the action they visualized in order to evaluate their level of expertise for each action, besides a general motor standardized evaluation. However, in line with other eye tracking studies in children and more particularly in ASD children, this study confirms that looking time and pupil dilation are interesting behavioral and physiological markers in autism. Further studies could make use of these markers in similar paradigms to evaluate the influence of motor experience during early interventions in very young children with ASD.

## Supporting information

Kernel maps of the 16 actions

## Acknowledgements

The authors wish to thank all the participants, children and families, for their valuable contribution. We also thank the two models who kindly accepted to be filmed for the videos. This work was supported by the Auvergne-Rhône-Alpes Academic Research Community (ARC2), and was performed within the framework of the LABEX CORTEX (ANR-10-LABX-0042) of Université de Lyon, within the program “Investissements d’Avenir” (ANR-11-IDEX-0007) operated by the French National Research Agency (ANR).

