## Supplementary figures and images for "Perception/action coupling in children with autism: insights from looking time and pupil dilation measurements"

### Kernel maps of the 16 actions

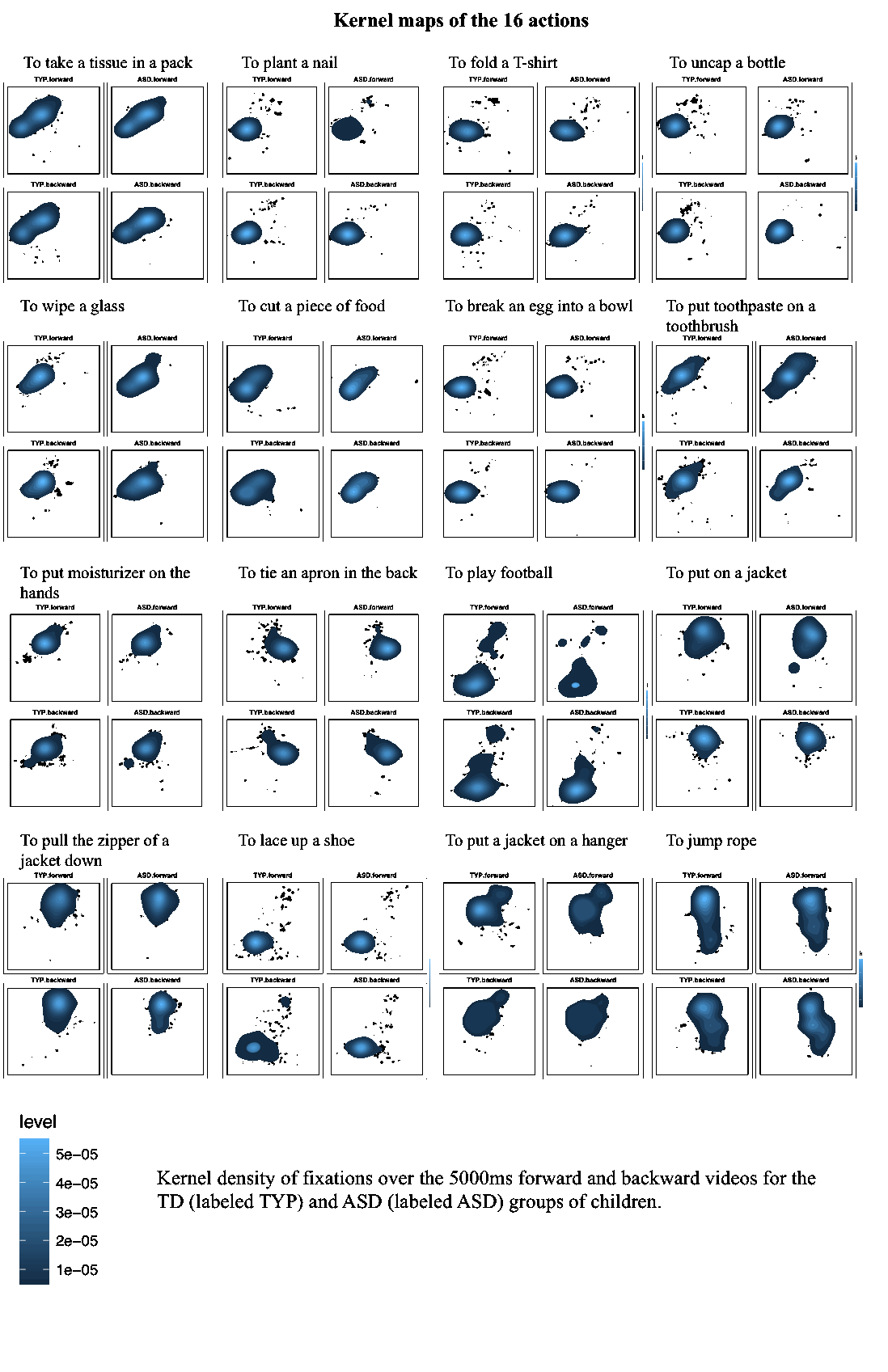
